# A multi-modal approach to decomposing standard neuropsychological test performance: Symbol Search

**DOI:** 10.1101/200998

**Authors:** Nicolas Langer, Erica J. Ho, Andreas Pedroni, Lindsay M. Alexander, Enitan T. Marcelle, Kenneth Schuster, Michael P. Milham, Simon P. Kelly

## Abstract

Neuropsychological test batteries provide normed assessments of cognitive performance across multiple functional domains. Although each test emphasizes a certain component of cognition, a poor score can reflect many possible processing deficits. Here we explore the use of simultaneous eye tracking and EEG to decompose test performance into interpretable, components of cognitive processing. We examine the specific case of Symbol Search, a “processing speed” subtest of the WISC, which involves searching for the presence of either of two target symbols among five search symbols. To characterize the signatures of effective performance of the test, we asked 26 healthy adults to perform a computerized version of it while recording continuous EEG and eye tracking. We first established basic gaze-shifting patterns in the task, such as more frequent and prolonged fixation of each target than each search symbol, and longer search symbol fixations and overall trial duration for target-absent trials. We then entered multiple such metrics into a least absolute shrinkage and selection operator (LASSO) analysis, which revealed that short trial completion times were mainly predicted by longer initial fixations on the targets and fewer subsequent confirmatory saccades directed back to the targets. Further, the tendency to make confirmatory saccades was associated with stronger gamma-amplitude modulation by mid-frontal theta-phase in the EEG during initial target symbol encoding. Taken together, these findings indicate that efficient Symbol Search performance depends more on effective memory encoding than on general “processing speed”.

## Introduction

In past decades, researchers have worked on accurately describing and understanding the various cognitive domains that are associated with mental health disorders. One important part of this effort has been to develop and apply neuropsychological test batteries that assess theoretically founded and usually comprehensive profiles of cognitive performance (e.g. Wechsler, Coalson, & Raiford, 2008). These tests have the advantage that the outcome measures typically hold high standards of test-quality (i.e. validity and reliability), and that individual outcome measures can be referenced to scores of a norm population. Thus, they offer objective quantifications of an individual’s performance across an extensive range of cognitive domains. However, performance in psychological tests usually reflects the product rather than processes of a cognitive function — or their dysfunction. As such, though psychological tests can highlight individual differences in aggregate cognitive performance, they say little regarding why. With the use of neuroimaging and electrophysiological techniques in cognitive neuroscience, new possibilities have opened up to gain insights into aspects of neural activity that correlate with such performance metrics. However, these insights are limited by the fact that each test, though designed to emphasize a certain component of cognition, call on multiple cognitive processes to support good performance, and poor scores could reflect a mixture of potentially heterogeneous processing deficits. Here, we report on the first step in a new initiative to decompose neuropsychological test scores into interpretable components of cognition through the use of eye tracking and scalp electroencephalography (EEG) recorded during test completion.

In the present study, we recorded EEG and eye tracking data during the performance of a computerized version of the “Symbol Search” task, a “Processing Speed” subtest of the Wechsler intelligence tests (Wechsler, 2008), which are widely used internationally for neuropsychological diagnostics. The Symbol Search subtest is designed to assess the speed and accuracy with which a subject can process nonverbal information (Wechsler et al., 2008). High scores require rapid and accurate processing of visual symbols that have no a priori meaning. The object of the test is to indicate whether either of two target symbols displayed on the left-hand side of each row is included in a “search set” of 5 symbols on the right-hand side, reporting simply ‘yes’ or ‘no’ for each trial. To do this a participant needs to (a) locate and encode the target symbols; (b) hold this information in short-term and/or working memory; (c) process each of the symbols in the search set, whether in turn or in parallel to some degree; (d) identify the symbol among the search set that matches one of the target symbols, or conclude that there is no match; (e) select and initiate the appropriate response (Joy, Kaplan, & Fein, 2004; Royer, Gilmore, & Gruhn, 1981). In routine clinical use, the overall score is calculated as the number of correct minus number of incorrect trials completed in a two-minute period, taken as an index of “processing speed.” This single aggregated score in itself offers no insight into whether poor performance arises, for example, from an inefficient active search strategy, deficient or rushed short-term memory encoding or faster memory decay, either overly hasty or overly deliberative policies for identifying each symbol, or general disengagement from the task. In principle, patterns of eye movements during the task should be indicative of the roles played by these different factors in overall test performance, and neural measures may provide convergent evidence. Although a previous fMRI study has highlighted brain regions that may participate in the task (Sweet et al., 2005), no study has yet parsed the more dynamic gaze-shifting patterns and concomitant electrophysiological processes during the test. In the present study we examined signatures of effective performance in adults, by quantifying measures such as fixation durations and frequencies and pupil size for each symbol within each relevant test subregion of the screen, alongside concurrently acquired neural activity. Our main goal was to conduct an exploratory analysis to pinpoint which EEG and eye tracking measures are most predictive of efficient performance of the task. Our results show that faster, correct trial completion was linked with eye movement behaviors reflecting stronger initial encoding of targets into memory, which was further supported by increased theta-gamma coupling (Axmacher et al., 2010; Sweeney-Reed et al., 2014) in the EEG.

## Methods

### Subjects

26 healthy young adults (age: 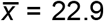, std = 7.01, females: 14) were recruited from the wider New York City-area community. According to the measures of subject handedness (Annett, 1970) all were right-handed. The study protocol was approved by the Institutional Review Board and was conducted in accordance with the latest version of the Declaration of Helsinki. All subjects provided written informed consent.

Prior to visiting the laboratory, participants completed a 10 min. pre-screening interview to confirm their eligibility and safety to participate in the study. This brief interview obtained information regarding an individual’s psychiatric history, including past or present diagnoses and/or treatment, as well as current medications and any neurological disorders. If a participant demonstrated no contraindications for EEG (e.g., history of seizures or epilepsy), he or she was then scheduled for a research study appointment.

### Task

The present study is a subproject of a recently launched initiatives to develop reliable dimensions of observable behavior and neurobiological measures for classification of mental disorders (in the spirit of RDoC of the NIMH) (Insel & Cuthbert, 2015). In this initiative, we have constructed a set of paradigms that provide objective, interpretable psychophysiological phenotypic markers in children and adults (Alexander et al., 2017). A subset of the data is publicly available (see Alexander et al., 2017; Langer et al., 2017). Here, we present the details of one particular paradigm, which is a computerized version of the *Symbol Search* subtest of the Wechsler Intelligence Scale for Children IV (WISC-IV), which together with the subtests Coding and Cancellation makes up the Process Speed Index (PSI) of the Wechsler Intelligence Scale for Children (WISC-IV) (Burgess, Flint, & Adshead, 1992; Lezak, 1995; Wechsler, 2004). We have obtained a research license agreement from Pearson to use the *Symbol Search* task for research purposes.

The present *Symbol Search* task is identical to the Wechsler Adult Intelligence Scale (WAIS-III) (Wechsler, 1997), which is equal to the Wechsler Intelligence Scales for children (WISC-IV) (Wechsler, 2004) and adults (WAIS-IV) (Wechsler, 2008) except the symbols are slightly different and presented in a different order. The Wechsler Intelligence Scales for children (WISC-IV) and adults (WAIS-IV) are the most commonly used intelligence batteries and frequently used in clinical routine (Rozencwajg, Schaeffer, & Lefebvre, 2010).

The visual geometric stimuli consisted of black symbols with a size of 1cm width and 1cm height (Figure 1). As on each page of the paper version, 15 trials were presented at a time on the screen in consecutive rows. Once a participant finished all 15 trials, they pressed the “next page” button to advance onward. There were 4 pages (a maximum of 60 trials) in total. No participant ever reached the end of the 60 trials. Each trial contained two target symbols and five search symbols, arranged horizontally across the row (Figure 1). Participants were instructed to indicate for each trial, whether either of the target symbols matched with any of the five search symbols. Responses were made by mouse-click and verified by the appearance of a cross over the checked ‘yes’ or ‘no’ checkbox. There were an approximately equal number of target-present (‘YES’) trials as target-absent (‘NO’) trials.

The participants had the option to correct their initial responses if they desired. As in the pen-and-paper clinical version, participants were instructed to solve as many trials as possible within two minutes. Before beginning the actual paradigm, participants performed a training block with 4 different trials not included in the main test, for which they received feedback, to ensure their comprehension of the task. No feedback was provided throughout the actual task.

### Data Acquisition

#### General

Participants were seated in a sound-attenuated and dark experiment room at a distance of 70 cm from a 17-inch CRT monitor (SONY Trinitron Multiscan G220, physical display dimension: 330 × 240 mm, resolution 800 × 600 pixels, vertical refresh rate of 100 Hz). A stable head position was ensured via the chin rest. Subjects were instructed to stay as still as possible during the tasks. Stimulus presentation was programmed in MATLAB (6.1, The Math-Works, Natick, MA, 2000), using the PsychToolbox extension (Brainard, 1997; Pelli, 1997). Instructions for the tasks were presented on the computer screen, and a research assistant answered questions from the participant from the adjacent control room through an intercom. The *Symbol Search* task was always performed following a 5 min resting EEG and 8 min sequence learning paradigm. Between each paradigm there was a break included and a new paradigm was started, when subjects have been ready to continue (see (Langer et al., 2017) for details).

#### Eye Tracking Acquisition

Eye position and pupil dilation were recorded with an infrared video-based eye tracker (iView-X Red-m, SMI GmbH; http://www.smivision.com/en.html) at a sampling rate of 120 Hz and an instrument spatial resolution of a nominal 0.1° and a gaze position accuracy of 0.5°. Prior to the start of task, gaze fixations were calibrated by asking the participant to focus their attention on each of 5 dots presented in a random order in either one of the 4 corners of the display space and in the middle of the screen. The calibration was repeated until the error between two measurements at any point was less than 2°, or the average error for all points was less than 1°.

#### EEG Acquisition

High-density EEG data were recorded at a sampling rate of 500 Hz with a bandpass of 0.1 to 100 Hz, using a 128-channel EEG Geodesic Hydrocel system (www.egi.com). The recording reference was at Cz (vertex of the head). The impedance of each electrode was checked prior to recording, to ensure good contact, and was kept below 40 kOhm.

### Behavioral Analysis

We analyzed the number of solved trials, errors, total score (number of correct trials - number of errors), accuracy, d-prime and criterion (Green & Swets, 1974; Macmillan & Creelman, 2005) to report the performance for each subject. The overall number of available trials for each trial type (YES, NO, correct and wrong, see below) is reported as well. Because of the high performance accuracy and low number of trials with errors, all subsequent eye tracking and EEG analyses focused on the trials, which were solved correctly.

### Eye Tracking Data Preprocessing

Saccades and fixations were detected with a dispersion based and a fixed-length moving interval algorithm. Since it is an SMI eye tracker we used, we chose to provide the fixation and saccade detections produced by SMI’s algorithm (BeGaze Software, version 3.5). For low speed eye tracking data (<200 Hz), it is recommended to choose dispersion based algorithms and the fixations as primary event (e.g. (Salvucci & Goldberg, 2000). Briefly, the algorithm identifies fixations as groups of consecutive points within a particular dispersion. It uses a moving window that spans consecutive data points checking for potential fixations. The moving window begins at the start of the protocol and initially spans a minimum number of points, determined by the given Minimum Fixation Duration (here: 50 ms) and sampling frequency. The algorithm then checks the dispersion of the points in the window by summing the differences between the points’ maximum and minimum x and y values and comparing that to the Maximum Dispersion Value; so if [max(x) − min(x)] + [max(y) − min(y)] > Maximum Dispersion Value, the window does not represent a fixation, and the window moves one point to the right. If the dispersion is below the Maximum Dispersion Value (here: 20 pixels), the window represents a fixation. In this case, the window is expanded to the right until the window’s dispersion is above threshold. The final window is registered as a fixation at the centroid of the window points with the given onset time and duration. Following this process, a saccade event is created between the newly and the previously created blink or fixation. A blink can be regarded as a special case of a fixation, where the pupil diameter is either zero or outside a dynamically computed valid pupil, or the horizontal and vertical gaze positions are zero. For the calculation of the pupil diameter, we used the bounding box method provided by SMI (BeGaze Software, version 3.5)^1^. In the bounding box method a squared bounding box is laid around the pupil. The pupil diameter is given in pixels as x and y values in of the bounding box.

### Eye Tracking Analysis

Our analysis strategy was to first provide a descriptive characterization of the gaze patterns within different subregions of the test for different trial types, and second, to systematically identify which measures were most strongly predictive of efficient performance.

#### Basic Eye Tracking Characteristics of Symbol Search Task

For each trial of the *Symbol Search* task all symbols and the check boxes for the responses were individually specified as regions of interest. The size of each region of interest was identical and defined as the size of the symbol (1cm^2^) plus horizontal extension of 0.75 cm on each side resulting in a region of interest size of 2.5x1cm per symbol or response check box (see Figure 1). No regions of interest were overlapping. Furthermore, the first two symbols were grouped to the label *targets*, the next five symbols were tagged as *search group* and the response check boxes were labeled as *responses* (Figure 1). For these specific sub-regions, we quantified pupil size, average fixation duration and number of saccade steps onto each symbol in the subregion (whether from a previous gaze location inside or outside the subregion). We quantified and compared these measures first across subregions, and second across trial types (‘YES’ Vs. ‘NO’ trials) within each subregion, initially without regard to “processing speed” performance (how fast the trial was completed).

**Figure 1.**
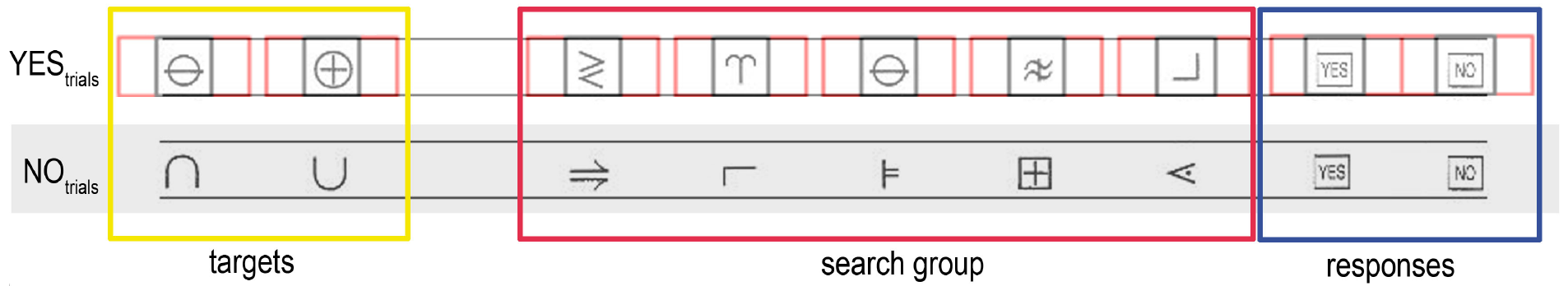
Graphic of Symbol Search task and region of interest for the eye tracker. Two representative trials of the Symbol Search task are displayed. For each trial all symbols and the check boxes for the responses were individually specified as regions of interest. The size of each region of interest (thin black box) plus horizontal extension on each side (thin red box) are indicated. The first two symbols were grouped to the label *targets* (yellow box), the next five symbols were tagged as search group (red box) and the response check boxes were labeled as responses (blue box).

For each of the three eye tracking measures (number of saccades per symbol, mean duration of fixation, and pupil size) we pooled the trials across subjects and conducted a hierarchical repeated-measures analysis of variance (rm-ANOVA), first of all with the factor of subregion (*targets, search group, response*) and a between-subject variable to adjust for between subjects differences. We then conducted a second rm-ANOVA for each eye tracking measure that instead considered the factor of trial type (YES vs. NO trials) within each subregion. We also analyzed the YES and NO trials in terms of differences in the total required time to solve the trial (trial duration). Significance level for the first and second analysis was set to p<0.05 (Bonferroni corrected; p = 0.05/3 = 0.017). Post-hoc paired sample t-tests were used to identify the specific differences.

#### Prediction of trial completion time based on Eye Tracking Measures

We next conducted a least absolute shrinkage and selection operator (LASSO) analysis to determine the eye tracking measures that best predicted how fast a subject can solve a trial in the *Symbol Search* task (trial duration) in order to gain insights into the mechanisms, strategies and behaviors associated with efficient performance of the task. We chose trial duration, to model each trial individually instead of an overall value, which enables more time sensitive analysis on a single trial level. For this analysis we normalized the eye tracking measures considered above: The number of saccades made to items in each subregion (targets, search set and response buttons) was normalized with respect to the total number of saccades in a trial (e.g. number of saccades on trials/total number of saccades). All measures of fixation duration measures were normalized (proportion of respective trial duration) by the total respective trial duration, and the pupil diameter measures were normalized (divided) by the pupil diameter at the start of the experiment. In addition to these basic eye tracking measures, we extracted other measures of specific interest. “*Confirmatory saccades on targets*” were identified as saccades made from any symbol in the search group back to the *targets* (with the precondition that the *targets* had been previously fixated). The “*1*^*st*^ *target fixation duration*” is quantified as the duration of the first fixation of the trial on the *target symbols*. We tested for differences between YES and NO trials regarding the number of confirmatory saccades made (Chi-square test) and the 1^st^ target fixation duration (t-test). On YES trials, we additionally extracted measures of gaze duration and number of fixations on the target-match, i.e., the symbol within the search group that matched one of the targets. Because of the different number of extracted measures and the significant differences between trial type (YES vs. NO trials: see result section and Figure 2), we computed the subsequent hierarchical LASSO analyses separately for the YES and NO trials to include all extracted measures for each trial type as potential predictors for the trial duration in the processing speed task. Before entering the analysis all eye tracking measures were z-transformed. All eye tracking measures are listed in Table 2 for the YES trials and Table 3 for the NO trials. The individual impact of these operationalized metrics on the individual trial duration was evaluated using a gradient ascent algorithm designed for generalized linear mixed models, which incorporates variable selection by L1-penalized estimation (glmmLasso.R) (Groll & Tutz, 2014). This is equivalent to a hierarchical LASSO analysis. Specifically, trial duration in the *Symbols Search* task was modeled considering a standard linear regression model for the observation of the response ***Y*** (Meier, van de Geer, & Buhlmann, 2008; Meinshausen & Yu, 2009):

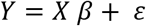

where ***Y*** was a vector of each trial duration. With 327 trials for YES conditions and 354 trials for NO condition, ***X*** was 327 by *27* for the YES and 354 by 16 for the NO trials (where 27 eye tracking measures were included in the LASSO analysis for YES trials and 16 eye tracking measures for the NO trials) and **ε** represents the noise vector with the zero mean and constant variance. As presented in (Tibshirani, 1996) the LASSO estimate is then given by:

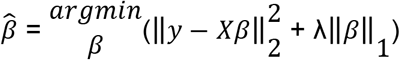

where ‖·‖_1_ and ‖·‖_2_ denote the *l*_1_-norm and *l*_2_-norm respectively. **λ** is a penalty parameter, which encourages a sparse solution 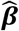 (potentially has entries equal to zero). The optimization problem is depicted by the equation above. The extended hierarchical version is described in (Groll & Tutz, 2014) and the associated R-package “glmmLasso” was used in the present analysis. The rationale to choose this hierarchical LASSO analysis is two fold: First, the high number of potential regressors. Unlike other methods such as multiple regressions, ridge regression or ordinary leasts quares, LASSO regression puts a sparsity constraint on 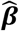 so that ***β*** can be zero and attempts to find the most informative eye tracker measures to predict trial duration (Tibshirani, 1996). The number of regressors selected by the LASSO operator is defined through the optimal tuning parameter **λ** with the lowest Bayesian information criterion (BIC, see (Groll & Tutz, 2014)). To determine the tuning parameter λ, we defined a fine grid of different values for the tuning parameter 0 ≤ λ_1_≤ … ≤ 1 (λ = 0-200). For each tuning parameter we calculated the BIC. Because of the hierarchical nature of the present lasso analysis, the typical lambda range from 0 to 1 is not appropriate here. We chose this specific range of λ, for which the highest lambda resulted in the suppression of all regressors to ensure all possible combinations. Next, the optimal tuning parameter was determined using the minimal BIC, and finally the whole data set is fitted again using the optimal λ parameter to obtain the final estimators (Groll & Tutz, 2014). Once the regressors were selected, we used bootstrapping to derive standard errors for the fixed effects estimates 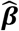 to estimate the significance of the individual regressors (see (Groll & Tutz, 2014) for details). Significance level was set to p<0.05.

## EEG Preprocessing & Analysis

### EEG Data Preprocessing

The non-scalp electrodes in the geodesics montage (chin and neck) were excluded, resulting in a standard 111-channel electrode array (e.g. Langer, von Bastian, Wirz, Oberauer, & Jancke, 2013). The EEG data were first high-pass filtered at 0.5 Hz and notch filtered (59-61Hz) with a Hamming windowed-sinc finite impulse response zero-phase filter (EEGLAB function pop_eegfiltnew.m). Bad electrodes were identified and replaced. Identification of bad electrodes was based on probability, kurtosis, and frequency spectrum distribution of all electrodes (Delorme & Makeig, 2004). A channel was defined as a bad electrode when recorded data from that electrode had a variance more than 3 standard deviations away from the mean across all other electrodes. This was realized with the eeglab MATLAB function: “pop_rejchan.m”. Subsequently bad electrodes were interpolated by using a using spherical spline interpolation (Perrin, Pernier, Bertrand, & Echallier, 1989) ‘eeg_interp.m’. Moreover, after automatic scanning, noisy channels were selected by visual inspection and interpolated. Contamination of eye movement was removed by linearly regressing 9 EOG channels from the scalp EEG channels (Wallstrom, Kass, Miller, Cohn, & Fox, 2004). Next, we used a robust Principal Components Analysis (PCA) algorithm, the inexact Augmented Lagrange Multipliers Method (ALM), to remove sparse noise from the data (see (Langer et al., 2017; Lin, Chen, & Ma, 2010 for details). The artifact-free EEG was recomputed against the average reference. All MATLAB codes for the preprocessing are implemented in the preprocessing toolbox “Automagic”, made by authors on this paper, and which is available at https://github.com/amirrezaw/automagic or as a plugin of EEGlab https://sccn.ucsd.edu/wiki/EEGLABExtensionsandplug-ins.

### EEG measures extraction

First the EEG and eye tracking data were synchronized using “EYE EEG extension” (Dimigen, Sommer, Hohlfeld, Jacobs, & Kliegl, 2011) to subsequently segment the EEG data based on the eye tracking measures. This approach enables EEG analysis to be carried out time-locked to the onsets of fixations (for an overview see (Dimigen et al., 2011)). The synchronization is performed in two steps. First, the eye tracking data were upsampled by linear interpolation to match the number of EEG sampling points. Afterwards “shared” events are identified. A linear function is fit to the shared event latencies in order to refine the estimation of the latency of the start- and end-event in the eye tracker recording. Synchronization quality was ensured, by comparing the trigger latencies recorded in the EEG and eye tracker data. All synchronization errors did not exceed 1 sample (2ms).

For the purposes of the current study, we were interested in oscillatory power in different frequency bands as well as cross-frequency phase-amplitude coupling (PAC), in particular theta-phase, gamma-amplitude coupling. PAC refers to the phenomenon in which dynamics of two frequency bands interact. PAC measures the degree to which the amplitude of a fast-frequency brain rhythm systematically depends on the phase of a slower-frequency rhythm (Cohen, 2017; Tort, Komorowski, Eichenbaum, & Kopell, 2010). In research on PAC in memory function, it has been proposed that the phase of a slower brain rhythm coordinates a temporal sequence of faster processes that represent specific items in memory or in sensory space (Axmacher et al., 2010; Canolty et al., 2006; Lisman & Idiart, 1995). In particular cross-frequency interactions between theta and gamma-band oscillations are a prominent feature of hippocampal but also cortical structures of the brain and thus could provide insights for encoding of multiple and sequentially ordered items intended to be memorized (Axmacher et al., 2010; Canolty et al., 2006; Tort, Komorowski, Manns, Kopell, & Eichenbaum, 2009). To compute PAC measures, we band-pass filtered the continuous EEG signals across the entire task period (including a 4sec. zero padding on the beginning and end of the task) for five different frequency bands resulting in a time-series for each frequency band. The independent frequency bands were determined following the classification proposed by Kubicki et al. (1979): theta (4-7.5 Hz), alpha1 (8-10 Hz), alpha2 (10.5-12.5 Hz), beta (12.5-30 Hz) and gamma (30.5-49.5 Hz). Equivalent results were found when the gamma band was defined between 30-80Hz. We then applied a Hilbert transform to each of these time-series resulting in a complex time-series,

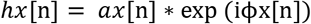

where *ax*[n]and iϕx[n] are the analytic amplitudes and phases, respectively, of a specific pass band *fp*[n].The phase time-series ϕx assumes values within (-π, π] radians with a cosine phase such that −/+π radians correspond to the troughs and 0 radians to the peak. The Hilbert phase and amplitude estimation method yields results equivalent to sliding window FFT and wavelet approaches (Bruns, 2004). We chose specifically the Hilbert transformation to maintain temporal information for the phase and amplitude of the frequency bands to enable PAC computation as well as the power of the different frequencies for time segments defined through fixations from the eye tracking recording. Theta-phase gamma-amplitude coupling was computed using an adapted version of the PACT eeglab plug-in (Miyakoshi et al., 2013) and the modulation index (MI) based on (Canolty et al., 2006) was used for all further analyses (Canolty et al., 2006; Cohen, 2017; Voytek, D’Esposito, Crone, & Knight, 2013) for a discussion of the PAC calculation).

### Mixed Linear Model (Statistical Analysis)

The EEG data were analyzed in response to the outcome of the hierarchical LASSO analysis on the eye tracking measures, which revealed that the “confirmatory saccades” (whether or not a confirmatory saccade was made to the targets) and the duration of the initial fixation on target are the best predictor for task performance. Thus, we hypothesized that the strength of the initial encoding of the target symbols determines if a confirmatory saccades is later conducted. In other words, the neural activity during the initial encoding of the targets predicts whether a subject performs a confirmatory saccade to the target again.

Thus we extracted the EEG activity during the 1^st^ fixation on the target symbols. Afterwards, we checked the segmented data for remaining artifacts and excluded trials if the amplitude of any EEG electrode exceeded ±90 μV. Bad trials were excluded from further analyses. The oscillatory power and the theta-phase gamma-amplitude coupling were extracted for each trial individually for the first fixation on the targets (average duration: 255.8 ms for YES trials and 269.9 for the NO trials). To test the statistical significance of the theta-phase gamma amplitude coupling, we performed a randomization analysis, in which we kept the actual amplitude and phase values at each trial, but randomized trial labels (following Voytek et al., 2013). This was done 1000 times. Thus we can obtain p-value of observing real theta-phase gamma-amplitude coupling given the null distribution of values obtained despite no systematic coupling. Subsequent analyses were only performed on electrodes, which displayed significant real theta-phase gamma-amplitude coupling (indicated in Figure 4). Moreover, we tested if the theta-phase gamma-amplitude coupling is contingent upon gamma amplitude, by including the gamma amplitude as an individual predictor into the subsequent statistical analysis.

EEG data were employed to predict, whether a confirmatory saccades was performed or not. Each trial was labeled as “trial with no confirmatory saccade” or “trial with confirmatory saccade”. We used an electrode-wise generalized linear mixed effects model (fitglme.m) with a logistic link function, because of the binary outcome variable: confirmatory saccades (Breslow & Clayton, 1993). The generalized linear mixed effects model is an extension to the generalized linear model, in which the linear predictor contains random effects in addition to the usual fixed effects. The fixed effects predictors were duration of the initial fixation on the target, trial order, the modulation index for the cross-frequency theta-phase gamma-amplitude coupling, as well as the oscillatory power of the five frequency bands during the time period the subject performed the first fixation on the target. A subject-wise random intercept was used to take between subjects differences into account. The significance level was set to p<0.05.

First, we performed the generalized linear mixed effects model including both trial types (YES and NO trials), for which we included the predictor (YES, NO). This is reasonable, because at the initial encoding participants don’t know if it’s a YES or NO trial and initial encoding should be indifferent for the two trial types. However, we additionally performed a mixed linear model for YES and NO trials independently to asses the specificity of the results within the different trial types. The trial type independent analyses revealed equivalent results as the combined analyses. All results are reported for the generalized linear mixed effects model, which included trial types as individual predictor.

## Results

### Behavioral Results

An overview of each subject’s performance and demographics is presented in Table 1. On average, subjects each solved 28 trials (sd: 5.69); average number of errors was 2.15 (sd: 2.17); mean accuracy was 92.14% (sd: 7.91); average d-prime 3.40 (sd: 0.95). Overall subjects solved 737 trials, of which 681 were correct and 61 incorrect. In more detail: all subjects together performed 366 YES trials (327 correct, 49 incorrect) and 371 No trials (354 correct, 31 incorrect). Because of the overall high accuracy on the low number of available trials with an error, all subsequent analyses were only performed for the correct trials.

**Table 1.** Overview of the subjects and the performance. For each subject the age, and the different performance measures are listed. In addition the average and standard deviation for each measures is specified.

|  | Age | Number of solver Trials | Number of Errors | Total Score | Accuracy | D-prime | Criterion |
| --- | --- | --- | --- | --- | --- | --- | --- |
| Subj 1 | 33 | 27 | 3 | 21 | 88.89 | 2.49 | 0.22 |
| Subj 2 | 25 | 30 | 2 | 26 | 93.33 | 3.44 | -2.44 |
| Subj 3 | 44 | 29 | 4 | 21 | 86.21 | 2.18 | -0.02 |
| Subj 4 | 38 | 23 | 1 | 21 | 95.65 | 3.66 | 2.33 |
| Subj 5 | 32 | 29 | 1 | 27 | 96.55 | 3.83 | 2.25 |
| Subj 6 | 24 | 26 | 2 | 22 | 92.31 | 3.35 | 2.49 |
| Subj 7 | 27 | 43 | 7 | 29 | 83.72 | 1.98 | -0.08 |
| Subj 8 | 27 | 40 | 1 | 38 | 97.50 | 3.97 | -2.18 |
| Subj 9 | 24 | 34 | 2 | 30 | 94.12 | 3.51 | 2.41 |
| Subj 10 | 19 | 23 | 6 | 11 | 73.91 | 2.21 | 3.06 |
| Subj 11 | 22 | 25 | 2 | 21 | 92.00 | 3.29 | 2.52 |
| Subj 12 | 23 | 33 | 0 | 33 | 100.00 | 4.65 | 0.00 |
| Subj 13 | 19 | 26 | 8 | 10 | 73.43 | 1.33 | 0.76 |
| Subj 14 | 22 | 29 | 1 | 27 | 96.43 | 3.83 | 2.25 |
| Subj 15 | 18 | 30 | 1 | 28 | 96.67 | 3.83 | -2.25 |
| Subj 16 | 18 | 28 | 1 | 26 | 96.15 | 3.79 | -2.27 |
| Subj 17 | 18 | 22 | 1 | 20 | 92.86 | 3.71 | -2.31 |
| Subj 18 | 18 | 28 | 4 | 20 | 85.13 | 2.89 | 2.72 |
| Subj 19 | 17 | 26 | 0 | 26 | 100.00 | 4.65 | 0.00 |
| Subj 20 | 18 | 21 | 0 | 21 | 100.00 | 4.65 | 0.00 |
| Subj 21 | 18 | 21 | 4 | 13 | 76.67 | 1.86 | 0.41 |
| Subj 22 | 17 | 35 | 2 | 31 | 95.56 | 3.16 | -0.01 |
| Subj 23 | 18 | 28 | 0 | 28 | 100.00 | 4.65 | 0.00 |
| Subj 24 | 18 | 26 | 2 | 22 | 90.91 | 2.85 | 0.00 |
| Subj 25 | 21 | 35 | 1 | 33 | 97.78 | 3.92 | 2.20 |
| Subj 26 | 17 | 20 | 0 | 20 | 100.00 | 4.65 | 0.00 |
| Average/ SD | 22.88<br>7.02 | 28.35<br>5.69 | 2.15<br>2.17 | 24.04<br>6.68 | 92.14<br>7.91 | 3.40<br>0.95 | 0.46<br>1.75 |

### Eye Tracking Analysis of Basic Characteristics of the Task

The repeated-measure ANOVA comparing the different regions of interest (*targets, search group, responses*) revealed significant differences between the number of saccades per symbol (F = 11.10; p = 0.0001). Post-hoc paired t-tests exhibited increased number of saccades on the *targets* compared to search group (t = 6.20, p = 1.75*10^−6^; the confidence interval for the difference (CI) = 0.279-0.56 fixations per symbol) and the *responses* (t = 11.34, p = 2.37*10^−11^; CI = 0.89-1.29 fixations per symbol). As would be expected, the number of saccades to *responses* were the lowest, significantly lower than the *search group* (t = 12.12, p = 5.80*10^−12^; CI = 0.56-0.79 fixations per symbol) (Figure 2A). To allow comparisons between regions of interest the number of saccades were calculated per symbol (2 symbols for *targets*; 5 symbols for *search group*; 2 symbols for *responses*) and relative to the total number of saccades (baseline correction).

The repeated-measures ANOVA for the duration of the fixations showed a significant effect for the within-subjects factor “region of interest” (F = 4.57, p = 0.015). The subsequent post-hoc t-tests demonstrated increased duration of the fixations on the *targets* compared to the *search group* (t = 5.51, p = 9.99*10^−6^; CI = 44.55-97.72 ms per symbol); and compared to the *responses* (t = 8.39, p = 9.68*10^−9^; CI = 103.33-170.54 ms per symbol). The average duration was also higher for the *search group* compared to the *responses* (t = 9.83, p = 4.55*10^−10^; CI = 52.01-79.59 ms per symbol). To enable comparisons between the different regions average duration was calculated per symbol and normalized to the respective trial duration. The ANOVA revealed no significant effect for the pupil size (F = 3.05; p = 0.16). All results of these analyses are summarized in Figure 2A.

In a subsequent analysis, we compared the NO (target not present in the search group) and YES trials (target present in the search group). The trial duration for the two trial types differed significantly (F = 6.02, p = 0.02). Subjects required significantly more time to solve NO trials compared to YES trials (t = 6.00, p = 2.86*10^−6^; CI = 0.68-1.40 seconds) (Figure 2B).

Investigating a possible difference between the numbers of saccades in NO trials compared to YES trials for targets, search group and responses revealed a significant main effect for trial type only for the *search group* (F = 8.41, p = 0.008). The post-hoc t-test found more saccades on the *search group* in the NO trials compared to YES trials (t = 8.21, p = 1.47*10^−8^; CI = 0.36-0.60 fixations per symbol). There were no significant differences between NO and YES trials regarding for number of saccades on the *targets* nor *responses.*

The comparisons between the NO and YES trials regarding the duration of fixations showed only for the *search group* a main effect for trial type (F = 7.59, p = 0.01). The post-hoc t-test revealed on average longer fixation durations on NO trials compared to YES trials on the *search group* symbols (t = 6.67, p = 5.52*10^−7^; CI = 34.83-65.97 ms per symbol). No differences between NO and YES trials were found for duration of fixations on the *targets* and *responses*. The comparison between the NO and YES trials for the pupil size revealed no significant differences for any region of interest. The comparisons between the YES and NO trials for the eye tracking measures are summarized in Figure 2C.

**Figure 2.**
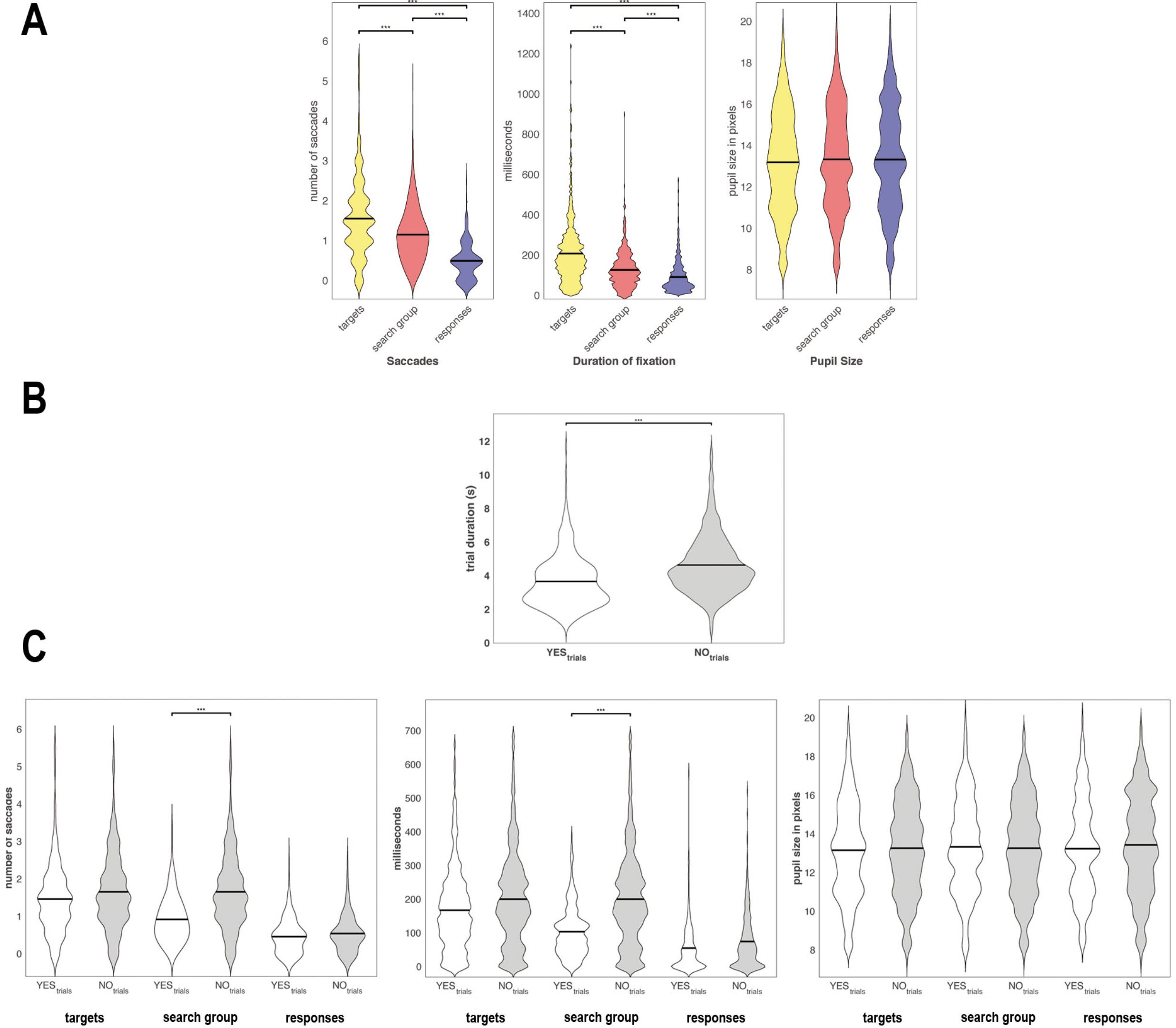
Basic Eye Tracking Characteristics of Symbol Search Task. In (A) the number of saccades per symbol (left), duration of the fixation (middle) and the pupil size (right) are displayed three different regions of interest. In (B) the different trial duration are depicted for YES (white) and NO (grey) trials. In (C) The different eye tracking measures as described in (A) are presented for YES and NO trials individually for each region of interest.

### Eye Tracking Lasso Analysis

To determine the eye tracking measures that best predicted processing speed performance in the *Symbol Search* task, all z-transformed eye tracking measures were entered into a hierarchical LASSO analysis. We performed for YES and NO trials an individually hierarchical LASSO analysis, because of the different amount of eye tracking metrics for the different trial types (more eye tracking measures for YES trials, see methods section).

In a fist step we determined the optimal tuning parameter λ. We tested a fine grid of different values for λ and calculated for each λ the corresponding BIC. The optimal λ was defined by the lowest BIC. Subsequently the whole data set was fitted again using the optimal λ (NO trials: λ = 151, BIC = -7577.2; YES trials: λ = 161, BIC = -4228.3) parameter to obtain the final estimators. The beta values of all eye tracking measures for the range of λ are plotted for YES and NO trials in Figure 3. For the NO trials the LASSO regression operator selected 4 of 16 eye tracking measures that significantly predicted trial duration for NO trials. Specifically, faster trial completion time was associated with (I) a lower total number of saccades onto targets ; (II) a lower number of confirmatory saccades onto targets; (III) a shorter proportion of the trial completion time spent gazing at the targets overall, without regard to when within the trial; but (IV) a longer proportion of the trial completion time dedicated specifically to the first, memory-encoding fixation on the targets. The coefficients estimates, z- and p-values for all regressors of the NO trials are depicted in Table 2. For the YES trials 6 of 27 eye tracking measures significantly predicted trial duration. Four of these were the same measures and directions as found for the NO trials (number of saccades on targets, occurrence of confirmatory saccades on target, duration of fixation on targets and 1^st^ fixation on the targets). In addition, the duration of the YES trials was also predicted by the occurrence of confirmatory saccades made specifically after fixation on the matching target, with longer completion time associated both with increased frequency and increased duration of confirmatory gaze shifts back to the target. The coefficients estimates, **z-** and p-values for all regressors of the YES trials are depicted in Table 3.

**Table 2.**
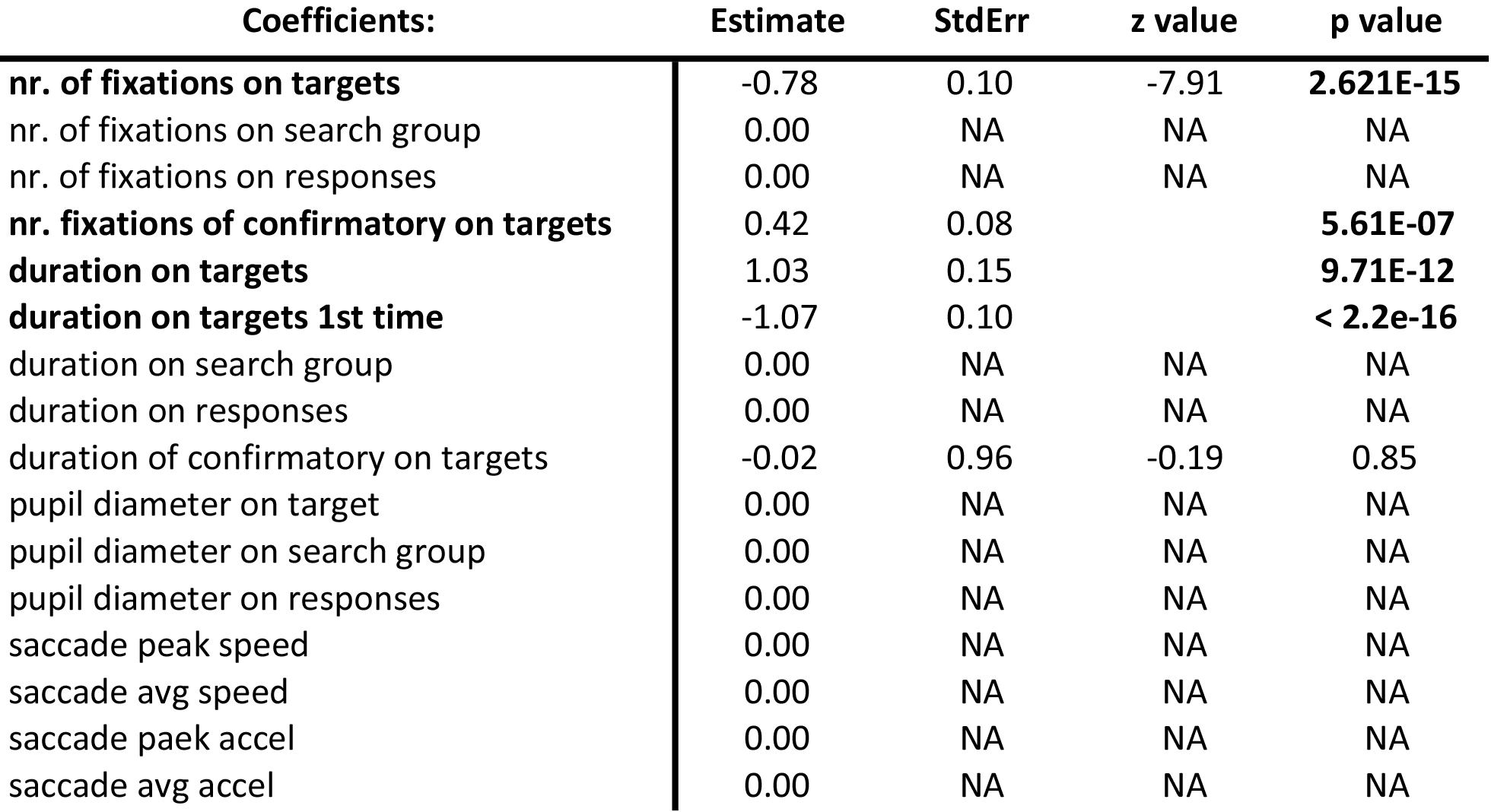
LASSO Coefficients for NO trials. All eye tracking measures for NO trials are listed. For each coefficient the beta estimate, the standard error, z- and p-value are presented

**Table 3.** LASSO Coefficients for YES trials. All eye tracking measures for YES trials are listed. For each coefficient the beta estimate, the standard error, z- and p-value are presented.

| Coefficients: | Estimate | StdErr | z value | p value |
| --- | --- | --- | --- | --- |
| **nr. of fixations on targets** | -0.91 | 0.15 | -6.04 | **1.522E-09** |
| nr. of fixations on search group | 9.54E+04 | 2.20E+05 | 0.43 | 0.66 |
| nr. of fixations on responses | 0.00 | NA | NA | NA |
| nr. of fixations on specific target | -0.13 | 0.15 | -0.88 | 0.38 |
| nr. of fixations on specific symbol in search group | -4.57E+04 | 1.05E+05 | -0.43 | 0.66 |
| nr. of fixations on other target | 0.00 | NA | NA | NA |
| nr. of fixations on other symbols in search group | -8.49E+04 | 1.95E+05 | -0.43 | 0.66 |
| **nr. fixations of confirmatory on targets** | 0.23 | 0.10 | 2.39 | **0.02** |
| **nr. fixations of confirmatory on specific target** | 0.60 | 0.16 | 3.71 | **0.0002** |
| **duration on targets** | 1.08 | 0.23 | 4.76 | **1.90E-06** |
| duration on search group | 1.01E+05 | 1.78E+05 | 0.57 | 0.57 |
| duration on responses | 0.00 | NA | NA | NA |
| **duration on targets 1st time** | -0.83 | 0.15 | -5.49 | **4.098E-08** |
| duration on specific target | 0.00 | NA | NA | NA |
| duration on specific target 1st time | -0.24 | 0.17 | -1.40 | 0.16 |
| duration on specific symbol in search group | -5.93E+04 | 1.04E+05 | -0.57 | 0.57 |
| duration on other target | 0.14 | 0.23 | 0.58 | 0.56 |
| duration on other symbols in search group | -9.01E+04 | 1.59E+05 | -0.57 | 0.57 |
| duration of confirmatory on targets | 0.19 | 0.15 | 1.25 | 0.21 |
| **duration of confirmatory on specific target** | -0.48 | 0.22 | -2.16 | **0.03** |
| pupil diameter on target | 0.00 | NA | NA | NA |
| pupil diameter on search group | 0.00 | NA | NA | NA |
| pupil diameter on responses | -0.07 | 0.06 | -1.17 | 0.24 |
| saccade peak speed | 0.00 | NA | NA | NA |
| saccade avg speed | 0.00 | NA | NA | NA |
| saccade peak accel | 0.00 | NA | NA | NA |
| saccade avg accel | 0.00 | NA | NA | NA |

**Figure 3.**
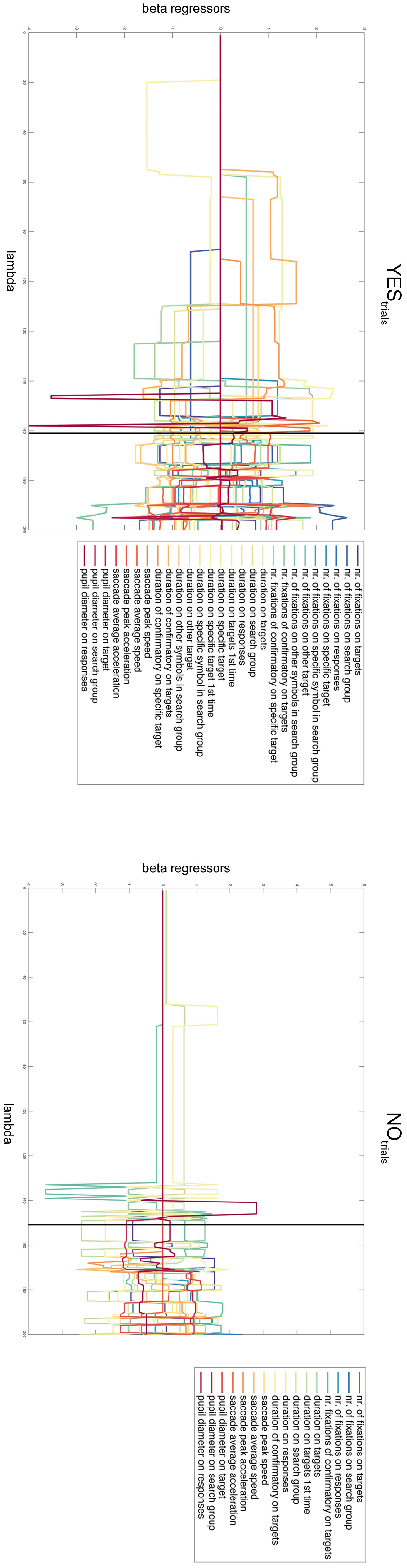
Hierarchical LASSO analyses of the eye tracker data. The results of hierarchical LASSO analyses of all eye tracking measures are presented for YES and NO trials individually. On the x-axis the different lambda values are outlined. The y-axis represents the beta regressor for each predictor at a specific Based on the results from the hierarchical LASSO analysis, we have performed post-hoc Chi-Square test to compare the proportion of number of trials with a confirmatory saccade to all trials between YES and NO trials. There was a trend that NO trials had a higher proportion of confirmatory saccades compared to YES trials (Chi2 = 3.08; p = 0.08). Moreover we conducted a t-test to compare the duration of the first fixation on the target symbols between YES (mean = 255.8ms, std = 224.1ms) and NO trials (mean = 269.9ms, std = 198.5ms). There were no differences between the YES and NO trials (t = -0.84; p = 0.40).

### EEG Results

The results of the LASSO analysis showed that faster trial completion time was associated with longer-duration first fixation on the targets and subsequently, fewer confirmatory saccades directed back to the targets, suggesting that the initial encoding of the targets into memory may be the most critical factor for efficient test performance. We investigated this further by analyzing signatures of memory encoding in the EEG activity during the initial fixation on the targets (REF). Based on the extracted EEG measures the goal was to predict a later occurrence of confirmatory saccades, by using a generalized linear mixed effects model with a logistic link function (binary outcome variable confirmatory saccades). We included and thus controlled for the following predictors: duration of the 1^st^ fixation on the target, trial type (YES or NO); trial order; the modulation index for the cross-frequency theta-phase gamma-amplitude coupling, as well as the oscillatory power of the five frequency bands during the first fixation on the target. The theta-phase gamma-amplitude coupling for the trials with and without confirmatory saccades are presented in Figure 4A. As specified by (Tort et al., 2010), we further divided the theta-phase into 10 equally sized bins (30 ° each bin) and the associated gamma amplitude for each bin are presented in Figure 4B.

The generalized linear mixed effects model revealed that mid-frontal theta-phase gamma amplitude coupling (modulation index) significantly predicts whether a later confirmatory saccades was conducted (Figure 4C), i.e., the higher the modulation index during encoding of the targets the less probable is the subsequent confirmatory saccade. There are further marginal effects in the oscillatory power of the theta frequency band in occipital and frontal electrodes (Figure 4D). We further conducted a generalized linear mixed effects model for YES and NO trials independently with the identical predictors. For both trial types mid frontal theta-phase gamma-amplitude coupling significantly predicted whether a confirmatory saccade was conducted.

**Figure 4.**
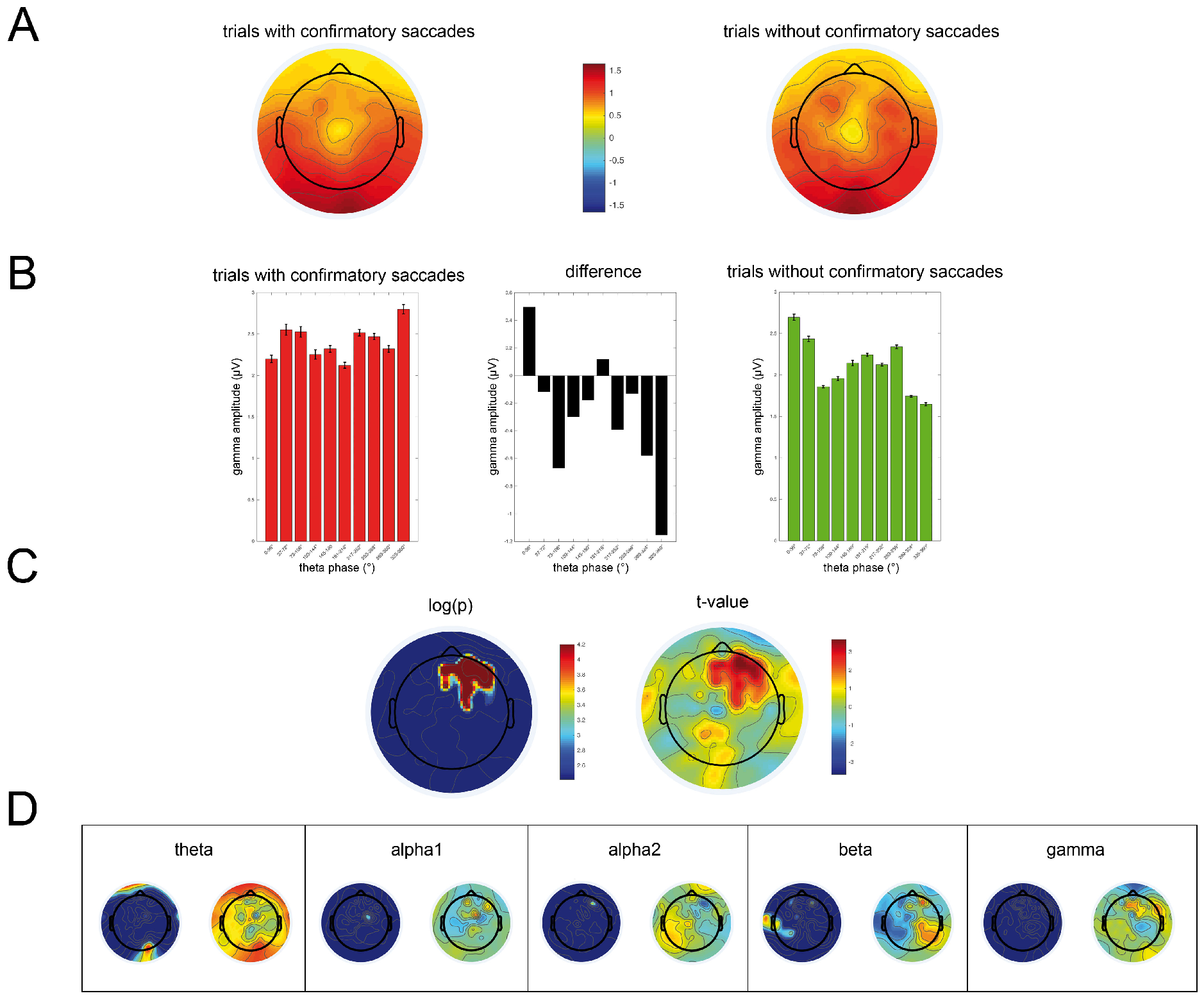
Results of the Logistic Regression of the EEG data. In (A) the theta-phase gamma-amplitude coupling during the initial fixation on the *target* is displayed for all electrodes and averaged across trials with confirmatory saccades (left) and trials without confirmatory saccades on *targets* (right). (B) Each bar represents a theta phase bins (bin width = 30°) and the corresponding gamma amplitude for trials with (red) and without (green) confirmatory saccades. In addition the difference between the two condition is displayed for each phase bin (black). (C) and (D) show the results of the logistic regression for the EEG data. (C) Mid-frontal theta-phase gamma-amplitude coupling significantly predicts if a subsequent confirmatory saccade was conducted. On the left side the log of the p-value is depicted and on the right side the t-value. The log of the p-values and the t-values for all other EEG measures is presented in (D) with the identical scale as in (C).

## Discussion

In the present study we described an investigation into using a multimodal eye tracking and EEG approach to decompose the aggregate performance metric furnished by the Symbol Search test into interpretable functional components. By analyzing basic gaze patterns among the relevant target, search group and response-button subregions, we first demonstrated that the subjects spent relatively more time processing each target symbol than each symbol in the search group, and also gazed longer at each symbol in the search group compared to the responses. It is important to note that these are relative measures, measured per symbol - since the *targets* subregion had two symbols and the search group consisted of five symbols, these differences were revealed despite the fact that cumulatively the subjects spent more total time gazing at symbols within the search group subregion than the target subregion. Furthermore, we showed that subjects needed more time to solve a NO trial compared to YES trials, which can be mainly attributed to the duration they spent in the search group. This result arises naturally from the fact that subjects can stop their search in the YES trials as soon as they have located the matching target, but must exhaustively check each search symbol in the NO trials.

Our LASSO analysis revealed that the occurrence of confirmatory saccades and the duration of the initial fixation on the *target* symbols were the main predictors for processing speed in the Symbol Search task. The longer the subject initially encoded the target symbols the faster they could solve the trial. At the same time, slower processing speed was observed when a subjects conducted confirmatory saccades on the target symbols. This can be explained by considering the most efficient course of task completion: the subject first encodes and memorizes the *targets*, then performs a visual search in the search group and subsequently chooses the appropriate response. However, if the subject’s encoding and memorization of the target symbols is not elaborated enough a confirmatory saccade on the *targets* is required to bolster or refresh the veracity of the memorized form of the target symbol, which leads to prolonged time to execute the task. One could thus speculate that longer duration of initial encoding leads to less confirmatory saccades, which then causes faster processing of the task, but an additional post-hoc analysis showed no difference between trials with confirmatory and no confirmatory saccades regarding duration of the initial fixation on the target symbols (t = 1.27; p = 0.21).

There are alternative explanations for confirmatory saccades, such as self-doubt, or second-guessing of the kind that is found in exacerbated form in obsessive-compulsive disorder (OCD). However, this seems to be unlikely in the present study, since only healthy subjects were included and our models account for subject-specific effects. This was substantiated by an additional post-hoc analysis of questionnaires regarding “uncertainty,” (CTAS) which could not identify any subjects with particular uncertainty (p > .42). However, personality traits (e.g. neuroticism, anxiety) promoting such behavior should be taken into account in studies with clinical populations. Taken together, these findings demonstrate that the initial encoding of the *targets* is crucial for fast performance of the Symbol Search task in healthy adults.

Concurrently acquired EEG during the initial fixation of the target symbols lended further support to the contention that time-wasting confirmatory saccades were a result of poor initial memory encoding of the targets. Specifically, significantly decreased mid-frontal theta-phase gamma-amplitude coupling and occipital and frontal theta power during target stimulus encoding were found in trials where at least one confirmatory saccade was made, compared to trials in which none were made.

Increased theta activity has been related to information encoding and retention in working memory tasks (Raghavachari et al., 2001; Raghavachari et al., 2006; Sauseng, Klimesch, Schabus, & Doppelmayr, 2005) and working memory training (Langer et al., 2013). Evidence from multiple studies indicates that theta is involved in timing coordinated activity within and across regions to enable successful encoding and retrieval (Johnson & Knight, 2015). The theta oscillations are considered as a result of an interaction within neuronal networks, mainly in the pyramidal cells of the hippocampus. Several feedback loops connect the hippocampal formation with different cortical regions, the prefrontal cortex in particular (Klimesch, 1999; Miller, 1991; Steriade, Jones, & Llinas, 1990). Fell et al. (2011) have demonstrated that successful encoding is associated with hippocampus coherence in the theta band and in the gamma band. Theta rhythms often do not occur in isolation but are frequently accompanied by activity in higher frequency ranges, in particular gamma frequencies (Haegens, Osipova, Oostenveld, & Jensen, 2010; Jokisch & Jensen, 2007; Palva, Kulashekhar, Hamalainen, & Palva, 2011; Roux & Uhlhaas, 2014). Cross-frequency coupling between theta and gamma frequencies has been linked to a variety of human cognitive processes, including learning (Sweeney-Reed et al., 2014; Tort et al., 2009), working memory performance (Axmacher et al., 2010; Roux & Uhlhaas, 2014), attention (Szczepanski et al., 2014) and reward processing (Cohen et al., 2009). Newer studies have repeatedly shown that cross-frequency coupling between theta and gamma may be used for information coding, if the lower frequency phase coordinates the activity of the subpopulations of cells that use higher frequency oscillations to process information (see Roux & Uhlhaas, 2014). Gamma band activity increases have been shown to represent individual stimuli in the neocortex and is sensitive to differences between stimuli (Rutishauser, Ross, Mamelak, & Schuman, 2010). These results suggest that the theta-phase gamma-amplitude coupling within and across regions may be central to human memory formation and capacity.

The fact that in the present study the mid-frontal theta phase gamma amplitude coupling predicts later confirmatory saccades, support the hypothesis of an imperfect initial encoding causing confirmatory saccades, which then results in a prolonged time to solve a task and consequently a decreased processing speed performance. Thus, this finding indicates that in healthy adults the performance in the Symbol Search task does not rely on “processing speed” characterized as a general brain capacity, but rather relies critically on specific cognitive components such as memory encoding. Future studies have to show whether our findings can be generalized to other demographic or clinical populations. Our newly assembled multi-modal test battery allows studying different possible deficits during the execution of the task. We expect that patients with different mental disorders exhibit problems in different aspects of the tasks, which might help to facilitate a more precise description of the deficits and identify possibly distinct subgroups within a given mental disorder. Because processing speed is a fundamental component of many cognitive functions, it may be particularly useful as a sensitive predictor of changes in higher-order cognitive abilities and an early marker of brain dysfunction (Duering et al., 2014; Eckert, 2011; T. A. Salthouse & Ferrer-Caja, 2003). Decreased processing speed has been associated with various psychiatric and neurological disorders (Donders, Tulsky, & Zhu, 2001; Duering et al., 2014; T. A. Salthouse & Ferrer-Caja, 2003; Wechsler, 2004), but also with decreased reading performance, intelligence (across all ages) (Fry & Hale, 2000; Nettelbeck & Young, 1989; Verhaegen, 2013) and increased mortality (Aichele, Rabbitt, & Ghisletta, 2015). But processing speed is also a prerequisite for everyday activities, for instance safe driving in old age (Vance, Heaton, Fazeli, & Ackerman, 2010). For this reason it is fundamental to adequately understand the measurements of processing speed used in psychiatry. In general the actual term ‘processing speed’ is often associated with aspects of measurement and there are a variety of different types of tasks used to assess processing speed. The variety of different measures for processing speed ranges from simple reaction time tasks to more complex tasks as included in the subtests of the WISC & WAIS. Nonetheless, previous studies have shown that subjects who are more efficient on one processing speed task tend to be more efficient on the others (Kuznetsova et al., 2016). As a result, in behavioral research data reduction techniques are often used, which have the advantage of removing error and task-specific variance associated with each individual test by combining an individuals performance on several tests into a general index of information processing speed (e.g. Kuznetsova et al., 2016; Penke et al., 2012). However as time and resources are limited in clinical practice, practitioners usually use the processing speed index of the WISC/WAIS, which is composed of the subtests Symbol Search, Coding and Cancellation. Little is known about the neural correlates of these tasks. Only one fMRI study investigated the Symbol Search task and has identified enhanced activity in bilateral medial occipital, parietal and dorsolateral prefrontal cortices (DLPFC) during the performance of the Symbol Search task compared to a control task (Sweet et al., 2005). Specifically, slower processing speed performance in the Symbol Search task was associated with increased activity in the DLPFC (Sweet et al., 2005). Whereas the study of Sweet et al. (2005) has provided new insights in the localization of the neural correlates of the Symbol Search task, a comprehensive view into the actual cognitive processes of the Symbol Search task, taking place at finer time scales, has been lacking up to now. The present study fills this gap, by exploiting simultaneous eye tracking and EEG measures, which offers temporally more detailed accounts of the underlying processing steps that lead to variability in performance of the Symbol Search task.

A wealth of neuroscientific studies has investigated the neural correlates of processing speed capacity, but a majority of studies have focused on related performance in very simple reaction time tasks or processing speed capacity as a general index. These studies consistently reported decreased processing speed to decreased white matter integrity (see T.A. Salthouse, 2017) for a review). While studies by Kuznetsova et al., (2016) and Penke et al. (2012) suggest rather global brain connectivity measures are correlated with processing speed, there has been mounting evidence for regional specificity of processing speed associations, such as a frontal network composed of ACC and DLPFC regions (Eckert, 2011; Takeuchi & Kawashima, 2012). In parallel processing speed has been mapped to neural activity. Functional neuroimaging evidence also implicates DLPFC involved with processing speed (Cabeza, 2002; Rypma, Berger, Genova, Rebbechi, & D’Esposito, 2005; Rypma & D’Esposito, 1999, 2000; Stebbins et al., 2002). Electrophysiological studies have identified a positive association between processing speed (measured with simple reaction time tasks), and P300 amplitude (Hansell et al., 2005) or the alpha frequency (see Klimesch, 1999; Klimesch, Doppelmayr, Schimke, & Pachinger, 1996) and reduced visual N100 amplitude (Wiegand et al., 2014).

Taken together the present study introduces a new approach to study the performance in the Symbol Search task. Using EEG and eye tracking enables objective and measurable insights in the underlying processes involved in this task, which allows to infer to the underlying etiology of low performance, which remains mainly unknown in the standard application of the Symbol Search task. Applied to a sample of healthy adults, we have identified that a deficient initial encoding of the target stimuli predicts succeeding confirmatory saccades, which is the main predictor for how fast a subject can solve the task. This finding suggests that the Symbol Search is not measuring processing speed per se, but rather memory encoding performance. Thus, this finding indicates that the successful execution of the Symbol Search task does require additional cognitive components, such as memory encoding.

## Footnotes

in mm not possible with this model of eye tracker

